# Structural analysis of the Sterile alpha motif (SAM) domain of the Arabidopsis mitochondrial tRNA import receptor

**DOI:** 10.1101/2023.11.19.566349

**Authors:** Bence Olasz, Luke Smithers, Genevieve L. Evans, Anandhi Anandan, Monika W. Murcha, Alice Vrielink

## Abstract

Mitochondria are membrane bound organelles of endosymbiotic origin with limited protein coding capacity. As a consequence, the continual import of nuclear-encoded protein and nucleic acids such as DNA and small non-coding RNA is required and essential for maintaining organelle mass, number and activity. As plant mitochondria do not encode all the necessary tRNA types required, the import of cytosolic tRNA is vital for organelle maintenance. Recently, two mitochondrial outer membrane proteins, named Tric1 and Tric2, for tRNA import component, were shown to be involved in the import of cytosolic tRNA. Tric1/2 binds tRNA^ala^ via conserved residues in the C-terminal Sterile Alpha Motif (SAM) domain. Here we report the X-ray crystal structure of the Tric1 SAM domain. We identified the ability of the SAM domain to form a helical superstructure with 6 SAM domains per helical turn and key amino acid residues responsible for its formation. We determined that the oligomerization of Tric1 SAM domain was essential for protein function whereby mutation of Gly241 resulted in the disruption of the oligomer and the loss of RNA binding capability in Tric1. Furthermore, complementation of *Arabidopsis thaliana* Tric1/2 knockout lines with a mutated Tric1 failed to restore the defective plant phenotype suggesting the oligomerization is essential for function *in planta*. AlphaFold2 structure prediction of the SAM domain and Tric1 support a cyclic hexamer generating a pore of sufficient dimensions to transfer tRNA across the mitochondrial membrane. Our results highlight the importance of oligomerization of Tric1 for protein function.

## Introduction

Mitochondria are essential organelles in many eukaryotic cells. The mitochondria arose over 1 billion years ago in an event where an archaebacterium and an eubacterium formed a symbiotic relationship (1). Due to the endosymbiotic origin, mitochondria maintain an active genome, encoding mitochondrial proteins and RNA products required for mitochondrial function. The number of active protein coding mRNA, tRNA and rRNA coding genes varies significantly between eukaryotes (2). However, much of the mitochondrial protein content is nuclear encoded and synthesized in the cytosol as preproteins. These preproteins are directed to the mitochondria for import via signal sequences, and imported into the mitochondrial intermembrane space via the TOM (translocase of the outer membrane) protein family and to the matrix via TIM (translocase of the inner membrane) proteins (3). The TOM and TIM complexes are involved mainly in protein import into the organelle. However, as some of the mitochondrial protein content is encoded in the organelle genome and not all RNA content is provided by the mitochondria, RNA import is essential for proper organelle function.

The mitochondrial genome encodes a number (varying between organisms) of genes to produce different tRNA species, however some tRNA is also imported from the cytosol. In some cases, the cytosolic tRNA imported is redundant with tRNA produced in the mitochondria. This redundancy may serve a regulatory function for protein production (4). In *Arabidopsis thaliana* (Arabidopsis), a plant specific mitochondrial tRNA import component has been identified (5). Named Tric1 and Tric2 (<u>tR</u>NA import <u>c</u>omponent 1 and <u>t</u>RNA import <u>c</u>omponent 2), these almost identical proteins belong to the <u>pr</u>eprotein and <u>a</u>mino acid <u>t</u>ransporters (PRAT) family of proteins but are distinct from other PRATs in that they also contain a C-terminal SAM (<u>s</u>terile-<u>a</u>lpha-<u>m</u>otif) domain (7).

The PRAT protein family is a large, conserved family of membrane proteins responsible for the transport of proteins and amino acids into both mitochondria and plastids (6-11). In *Arabidopsis*, ten of the sixteen members of the PRAT family are found in the mitochondrial inner membrane and include the protein import transporters Tim17, Tim22, and Tim23 (7). Another three, including OEP16, are located in the chloroplast outer envelope and have been shown to be involved in amino acid transport (7). Tric1 and Tric2, have been shown to be dual-targeted to both the mitochondria and chloroplast and, in mitochondria, they have been implicated in tRNA import. This is facilitated by the SAM domain, which is directly involved in tRNA binding (5).

SAM domains, first characterized in yeast and *Drosophila melanogaster* (12) are composed of ∼65-70 amino acids and participate in diverse developmental processes including sexual differentiation in yeast and spatial regulation during embryonic development in *Drosophila melanogaster* (13). Since the discovery of this domain, it has been identified in numerous diverse proteins in a wide range of eukaryotes (14–16), including plants (17,18), and even in some prokaryotes (19).

The SAM domain was originally described as a protein-binding domain, involved in protein-protein interactions. They have been observed in head-to-tail homo-SAM interactions (20) or hetero-SAM interactions (21,22) forming linear oligomeric structures of differing stoichiometries (23). SAM domains have also been observed in heterotypic protein interactions such as shown in Ets2, a human transcription factor, binding Cdk10 (cyclin dependent kinase 10) with the interaction implicated in transactivation (24) or BAR (bifunctional apoptosis regulator) interacting with Bcl-2 (B-cell lymphoma-2) and Bcl-X_L_ (B-cell lymphoma-extra large), where they play a role in regulating apoptosis (25).

In addition to protein-protein interactions, SAM domains have been observed to interact with RNA. The first SAM domain containing protein shown to bind RNA was Smaug from *Drosophila melanogaster* (26,27). The structure of Vts1, a yeast RNA binding protein and homolog of Smaug, has been solved by NMR spectroscopy (28), describing the RNA binding surface of the SAM domain and identifying key residues required for RNA interaction. In *Arabidopsis,* Tric1 and Tric2 SAM domains have been shown to bind tRNA^ala^ via key lysine residues at positions 205 and 210, and are required for tRNA import into the mitochondria *in vivo* (5).

Here we describe the crystal structure of the SAM domain from Tric1. We show it has the ability to oligomerize to form a helical superstructure with 6 SAM monomers per helical pitch and identify key residues required for this oligomerization. *In vitro* RNA binding assays reveal these residues are also required for RNA binding and complementation of the double Tric1 and Tric2 knockout mutant line (*tric1:tric2)*. Finally, modelling with AlphaFold of the full length Tric1 protein supports the hexameric form, hinted to in the crystal structure.

## Results

### The crystal structure of WT Tric1 SAM domain

The crystal structures of the Tric1 SAM domain (residues 191 – 261) as well as two variants (Asp235Ala and Gly241Glu) have been determined. The wildtype (WT) SAM domain structure was determined by MAD phasing from a selenomethionine mutant; the protein chain has three methionine residues. The final structural model was refined to 1.48 Å resolution. The asymmetric unit of the crystal for the WT structure contains three monomers with identical contacts between the three protein chains. The protein adopts a characteristic fold consisting of a 5-alpha-helical bundle (Fig. 1A). The overall globular structure consists of four short helices (H1-4) and one long helix (H5) at the C-terminal end of the protein. Comparison to other SAM domains (16,17,26,28) indicates that the overall fold of the protein and secondary structure elements are highly conserved, and that the Tric1 SAM domain has slightly more loop regions and shorter helices (Fig. S1A). The observed electron density for the entire protein chain is well defined, with the exception of three positively charged residues at the C-terminus (Lys259, Arg260 and Lys261). This structural conservation between SAM domains in different proteins is common despite low sequence conservation (Fig. S1B). Low sequence identity is observed between SAM domains involved in RNA binding (Tric1, Vts1, and Smaug) and those not involved in RNA binding (LFY and PHC3) (Fig. S1B).

**Figure 1:**
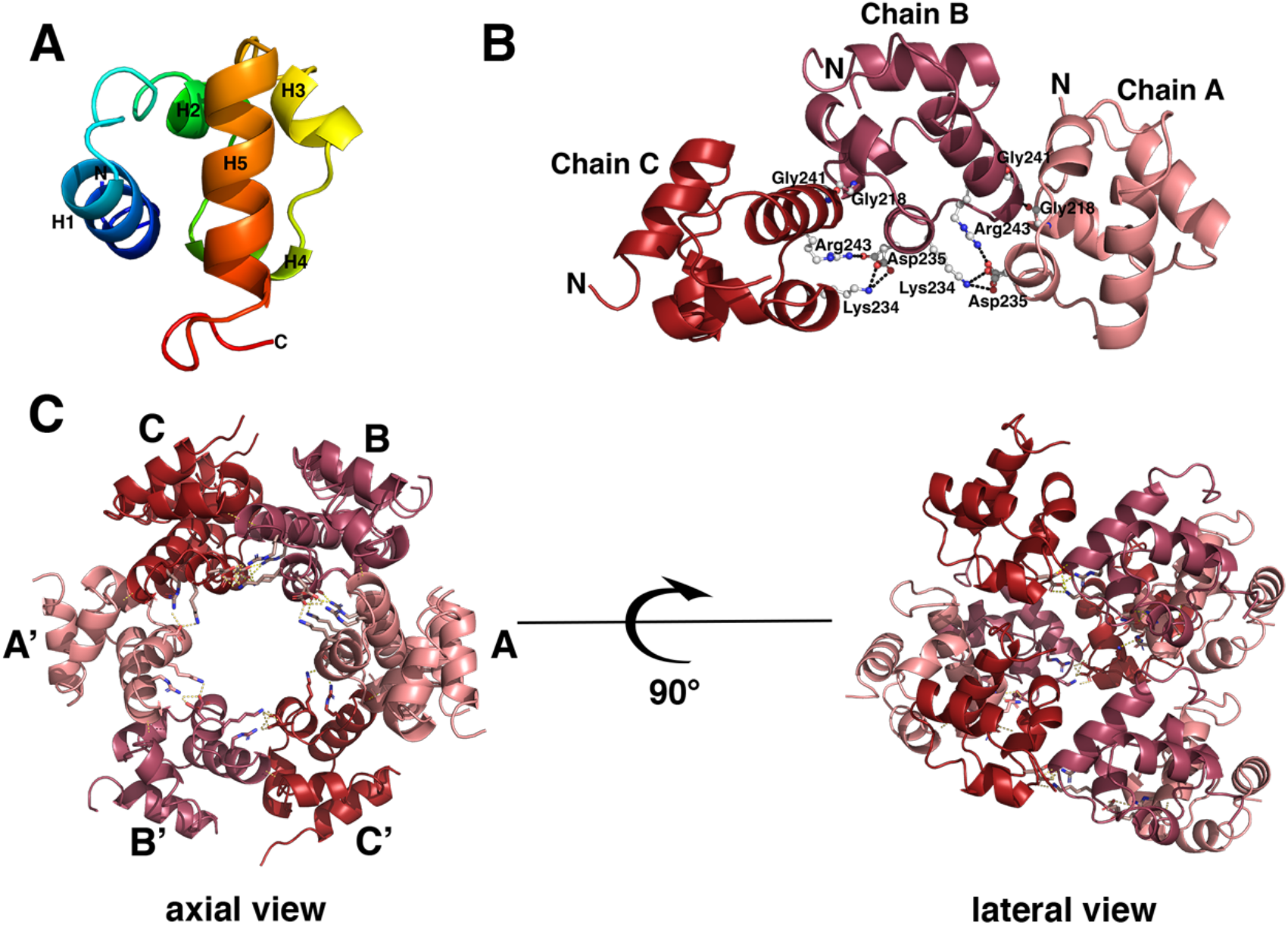
The structure of the *Arabidopsis thaliana* Tric1 SAM domain. **A.** A ribbon representation of the monomer showing the secondary structure elements. The protein is colored from blue for the N terminus to red for the C terminus with the five alpha helices labelled. **B.** The intermolecular interactions between the monomers of the asymmetric unit in the WT structure. Each monomer is shown in cartoon representation and colored different shades of red. The monomers are labelled as Chain A, B and C. The residues involved in the inter-subunit interactions are shown in stick representation with the hydrogen bonds displayed as dashed lines. **C.** The superstructure formed by the WT Tric1 SAM domain corresponding to three asymmetric units of in the crystal lattice shown from an axial and a lateral view. The monomers in one asymmetric unit are labelled as A, B and C and the corresponding monomers in the second asymmetric unit are labelled as A’, B’ and C’. The third asymmetric unit monomers lie directly below those of the first asymmetric unit.

The oligomeric structure within the asymmetric unit shows a number of side chains that are involved in hydrogen bond and salt bridge contacts between the subunits (Fig. 1B). Further analysis of the crystal lattice revealed a helical superstructure, similar to other SAM domains (16,20,29-31). This superhelix consists of six monomers per turn with a pitch of 28 Å (Fig. 1C). An electrostatic surface of the superhelix reveals a positively charged surface along one face of the six-subunit structure near the amino end of the SAM domain and a negatively charged surface along the opposite face of the structure near the carboxyl end of the SAM domain (Fig. S2). This charge complementarity likely contributes towards the lattice packing. It is noteworthy that residues prior to the amino end of the SAM domain in the full-length protein may affect the electrostatics of this surface. Intermolecular subunit interactions are more extensive (397 Å^2^ buried surface) between adjacent molecules laterally than between molecules along the pitch of the superhelix (251 Å^2^ buried surface). Besides the lateral interactions, intermolecular interactions between monomers along the helical axis further strengthen the superstructure.

To assess whether the oligomeric structure observed in the crystal structure is also present in solution, pure WT SAM domain was eluted from a Superdex75 gel filtration column and the peaks analysed by Western Blot to confirm the size of the protein (Fig. S3). Pure SAM domain eluted from the column as two main species, one at a retention volume of ∼58 ml representing an oligomeric species of the size equal to the BSA standard of 66kDa, and a further peak at a retention volume of ∼90 ml representing the monomeric SAM domain (Fig. S3a). Samples from each of the peaks were heat treated or left unheated prior to loading on an SDS gel followed by Western Blot analysis using a monoclonal anti-polyhistidine-peroxidase antibody to identify specifically the histidine tagged version of the SAM domain in each of the peaks. Interestingly, both the heated and unheated samples derived from the sample that eluted at ∼58 ml appeared on the blot at a high molecular weight than would be expected for the monomer (∼11 kDa) suggesting a highly thermally stable and folded oligomeric form of SAM domain in solution. The band corresponding to the oligomeric species appears at approximately 50 kDa on the blot however, given that the protein is folded (as suggested by oligomerisation) it is likely to run as a smaller size than expected for a completely unfolded protein. This result provides evidence that the SAM domain forms a thermally stable oligomeric species.

### Gly241 is required for the formation of Tric1 SAM domain helical superstructures

To determine if the homo-oligomerization ability of the Tric1 SAM domain is required for protein function, mutational studies were undertaken. Based on the structural analysis of Tric1 SAM domain and the interactions between monomers in the helical superstructure, several mutants were designed. Firstly, Asp235Ala, which would eliminate the salt bridge interaction between the carboxylate side chain of Asp235 and the side chains of Lys234 and Arg243 of a neighbouring monomer, and Gly241Glu, which is hypothesised to push two neighbouring monomers away from one another due to the extended glutamate side chain, as well as introducing a negative charge.

The crystal structure of the Tric1 SAM Asp235Ala mutant was determined and refined to 2.07 Å resolution revealing an identical fold and lattice packing to that of the wild-type (WT) protein (Fig. S4A, B). As expected, the salt bridge interaction between position 235 and Lys234 and Arg243 is absent; hydrogen bond interactions are still present between the main chain oxygen of Gly218 of monomer A and the main chain nitrogen of Gly241 of monomer B (Fig. 2A). Additionally, the mutant adopted an identical superhelical structure to that observed for the Tric1 SAM domain (Fig. S4A and B).

**Figure 2:**
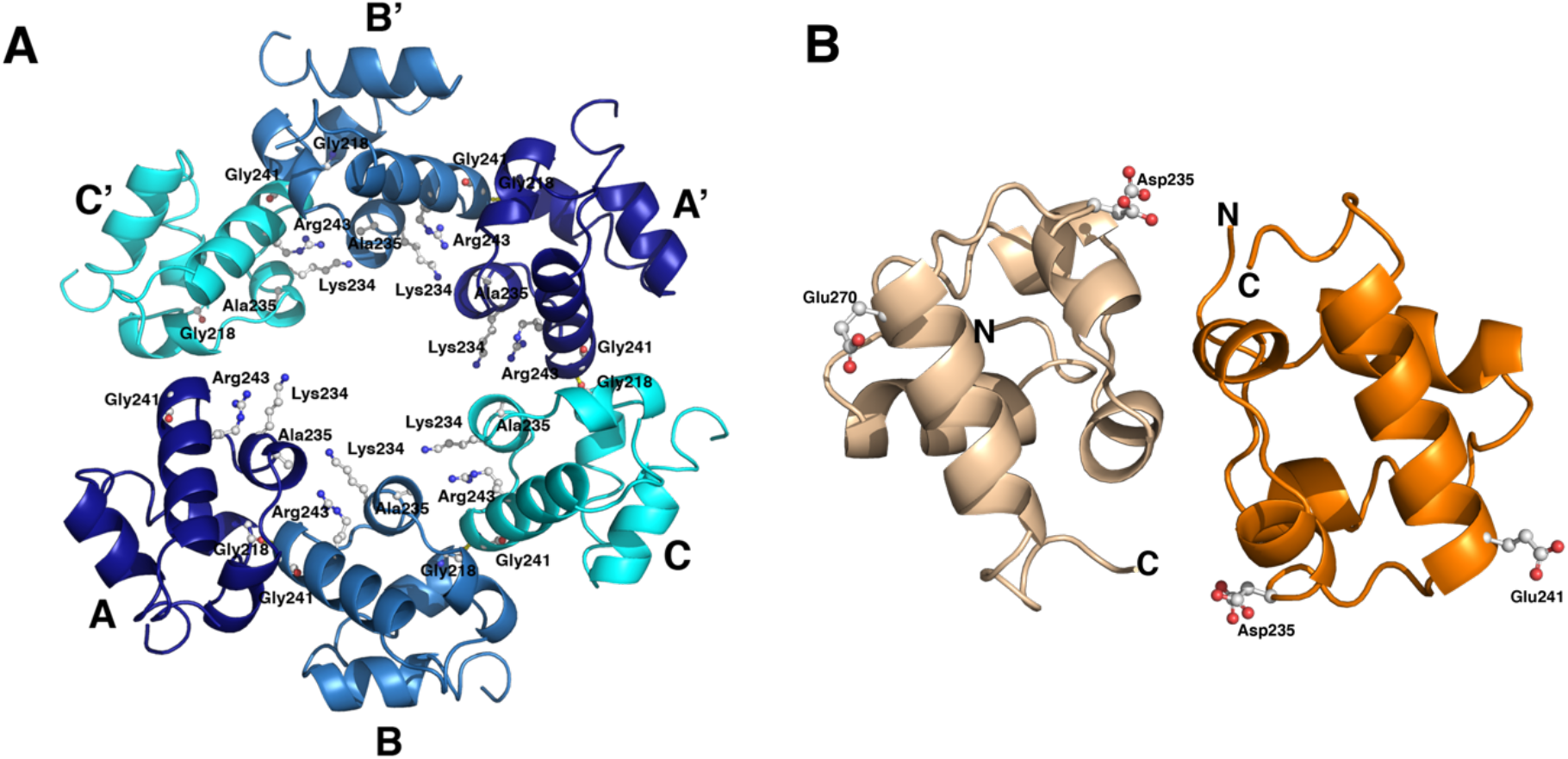
Structures of the SAM domain variant proteins. **A.** The Asp235Ala variant structure showing two asymmetric units corresponding to 6 monomers. The monomers in one asymmetric unit are labelled as A, B and C and those in the second asymmetric unit are labelled as A’, B’ and C’. The monomers are shown as a ribbon representation in different shades of blue. **B.** The non-crystallographic symmetry dimer in the asymmetric unit of the Gly241Glu structure. The N and C termini for each monomer in the Gly241Glu structure are labelled. Each monomer is shown in a different shade of orange. Individual amino acid side chains are shown in ball and stick representation. The side chain corresponding to Asp235 is shown in two conformations.

Crystals of the Gly241Glu mutant were obtained but from different crystallization conditions suggesting a different packing to the WT SAM domain and to the Asp235Ala mutant. Furthermore, despite the identical space group for all three structures, the unit cell dimensions of the Gly241Glu mutant crystal structure differed from those of the WT Tric1 SAM domain and the Asp235Ala mutant (Table 1). The structure was determined and refined to 1.89 Å resolution. While the overall fold of the monomer does not differ from that of the WT protein or the Asp235Ala mutant, the number of molecules in the asymmetric unit and the overall crystal packing did differ significantly. Two molecules are present in the asymmetric unit and the polar intermolecular contacts between the two monomers was notably absent (Fig. 2B). Furthermore, the crystal lattice no longer showed a helical superstructure (Fig. S4C). Thus, the introduction of a longer side chain at position 241 in the amino acid sequence was sufficient to eliminate the subunit arrangement of the structure in the crystal lattice.

**Table 1:** Crystallographic data collection and refinement statistics.

| Protein | Se-Met |  |  |  | WT | Asp235Ala | Gly241Glu |
| --- | --- | --- | --- | --- | --- | --- | --- |
|  | Inflection Point | Peak | Low Remote | High Remote |  |  |  |
| Beamline | AS MX1 | AS MX1 | AS MX1 | AS MX1 | AS MX1 | AS MX1 | AS MX2 |
| *Data collection statistics* |  |  |  |  |  |  |  |
| Space group | $P2_12_12_1$ | $P2_12_12_1$ | $P2_12_12_1$ | $P2_12_12_1$ | $P2_12_12_1$ | $P2_12_12_1$ | $P2_12_12_1$ |
| Unit cell dimensions (Å) | | | | | $a=28.6$ | $a=28.7$ | $a=46.6$ |
| | | | | | $b=92.1$ | $b=92.6$ | $b=58.5$ |
| | | | | | $c=93.3$ | $c=94.1$ | $c=61.9$ |
| Resolution (Å)* |  |  |  |  | 1.48<br>(1.53-1.48) | 2.07<br>(2.14-2.07) | 1.89<br>(1.96-1.89) |
| Total reflections | 99,311 | 99,689 | 100,670 | 16,1549 | 599,785<br>(52161) | 220,700<br>(20864) | 183,488<br>(18001) |
| Unique reflections | 20,597 | 20,608 | 20,709 | 35,642 | 42,131<br>(4,135) | 15,930<br>(1,505) | 14,037<br>(1,355) |
| Completeness (%) |  |  |  |  | 99.9<br>(99.9) | 97.3<br>(97.7) | 99.5<br>(96.1) |
| $R_{merge}$ | | | | | 0.05<br>(1.20) | 0.14<br>(0.72) | 0.09<br>(1.50) |
| $R_{pim}$ | | | | | 0.01<br>(0.34) | 0.03<br>(0.019) | 0.02<br>(0.42) |
| Average I/$\sigma$ | | | | | 36.3 (2.1) | 19.5 (4.0) | 18.9 (2.4) |
| Multiplicity |  |  |  |  | 14.2 (12.6) | 13.9 (13.9) | 13.1 (13.1) |
| CC$_{1/2}$ | | | | | 1 (0.75) | 1.0 (0.91) | 1.0 (0.78) |
| *Refinement statistics* |  |  |  |  |  |  |  |
| $R_{work}$ | | | | | 0.18 (0.25) | 0.25 (0.28) | 0.20 (0.29) |
| $R_{free}$ | | | | | 0.20 (0.28) | 0.27 (0.34) | 0.24 (0.31) |
| # Protein residues |  |  |  |  | 198 | 193 | 125 |
| # Solvent molecules |  |  |  |  | 180 | 226 | 81 |
| RMS (bonds) (Å) |  |  |  |  | 0.012 | 0.005 | 0.004 |
| RMS (angles) (°) |  |  |  |  | 1.2 | 0.8 | 0.8 |
| Ramachandran favoured (%) |  |  |  |  | 99.5 | 98.9 | 97.5 |
| Ramachandran outliers (%) |  |  |  |  | 0 | 0 | 0 |
| Average B-factors (Å<sup>2</sup>) |  |  |  |  | 29.3 | 26.1 | 41.78 |

\*Values in parentheses are for the highest resolution shell.

### Asp235 and Gly241 are required for SAM domain RNA binding ability

It was previously shown that recombinant Tric1 SAM domain (residues 191 - 261) binds the T-arm of tRNA^ala^ via conserved lysine residues at positions 205 and 210 (numbered as Lys 15 and Lys20 of the SAM domain alone) (5). To determine if the oligomerization ability of Tric1 SAM domain was required for this binding, the RNA binding ability of Tric1 SAM Asp235Ala and Gly241Glu variants was tested. Electrophoretic mobility shift assays (EMSA) using fluorescein-labelled T-arm oligonucleotides from tRNA^ala^ were carried out (Fig. 3). Incubation of Tric 1 SAM domain with tRNA^ala^ resulted in a shifted band indicating RNA binding, as previously observed (5). No mobility shift bands were observed for the Tric1 SAM variants Asp235Ala, Gly241Glu or for the double variant Asp235Ala:Gly241Glu suggesting the abolishment of the tRNA binding capacity (Fig. 3A). Coomassie staining confirms purity and equal concentration of recombinant protein used in the assay (Fig. 3B)

**Figure 3:**
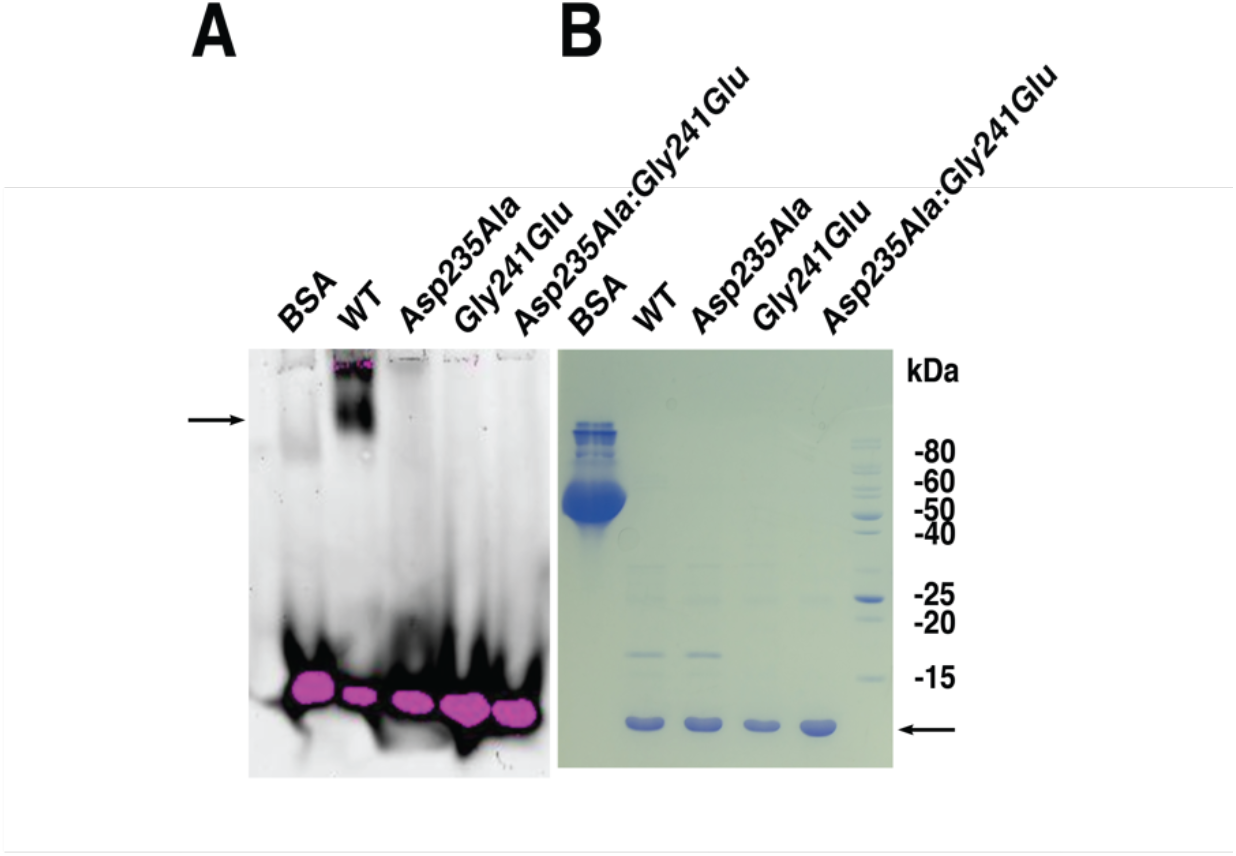
RNA Electrophoretic mobility shift assays of Tric1 SAM domain. **A.** WT Tric1 SAM domain and the variants Glu70Glu, Gly241Glu and Glu70Glu:Gly241Glu were electrophoresed with a fluorescein-labelled T-arm of tRNA^ala^. The arrow indicates the position of the WT Tric1 SAM domain shifted band. **B.** All the protein samples used in the electrophoretic mobility shift assay were resolved using 15% SDS-PAGE and Coomassie stained. The arrow indicates the position of the purified Tric1 SAM domain protein and variants.

### Asp235 and Gly241 are required for Tric1 protein function in planta

Biochemical assays suggest that the mutations studied abolish tRNA binding, potentially by disrupting oligomerisation (in the case of Gly241Glu). (Fig. 3). To determine if these residues impact Tric function *in planta,* complementation assays were carried out on the *tric1:tric2* knockout line using the full length WT Tric1, and Asp235Ala:Gly241Glu variants. The *tric1:tric2* knockout exhibits a distinct small, pale and developmentally delayed phenotype, which could be restored by complementation with WT Tric1. This complementation assay was repeated and shown to again restore the defective phenotype (Fig. 4A) (5). Independent complemented lines: WT 1 and WT 2, exhibited larger plants than the mutants with significantly increased chlorophyll content (Fig. 4A and B). Complementation of *tric1:tric2* knockout with the double mutant Asp235Ala:Gly241Glu failed to restore the WT phenotype (Fig. 4A and B). Two independent complementation lines: Asp235Ala:Gly241Glu exhibited the small developmentally delayed growth phenotype with significantly decreased chlorophyll content compared to WT complementation lines (Fig. 4B). These results suggest that Asp235 and Gly241 are required for functional complementation *in planta*.

**Figure 4:**
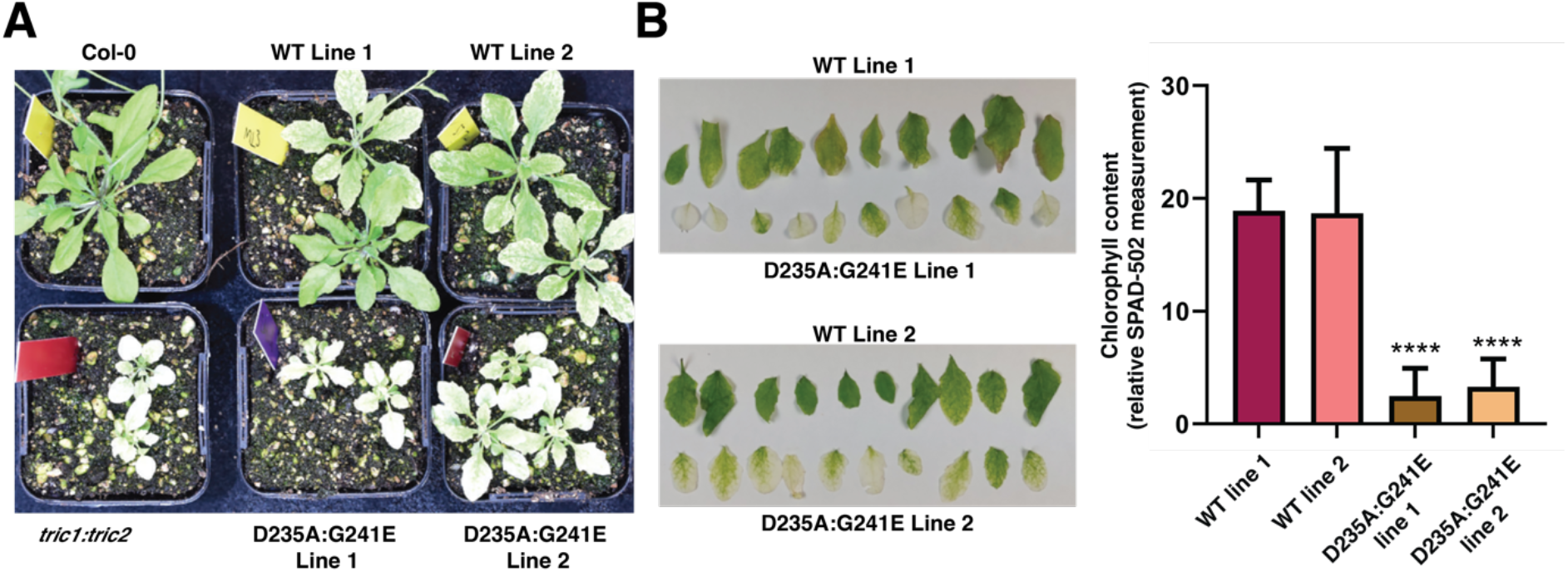
The effect of key amino acid mutations on Tric1 function *in vivo*. **A.** Complementation of the *Arabidopsis thaliana tric1:tric2* mutant with the full-length Tric1 cDNA (WT lines 1 & 2) and the Tric1 Asp235Ala:Gly241Glu double mutant construct (D235A:G241E lines 1 & 2). **B.** Chlorophyl measurements of the complemented plant lines. Each bar shows the average data from 5 technical replicates across one leaf from ten individual plants. Significant differences are indicated by **** (±sd, P < 0.0001, one way ANOVA from the Brown-Forsythe and Welsch tests, n = 10).

**Figure 5:**
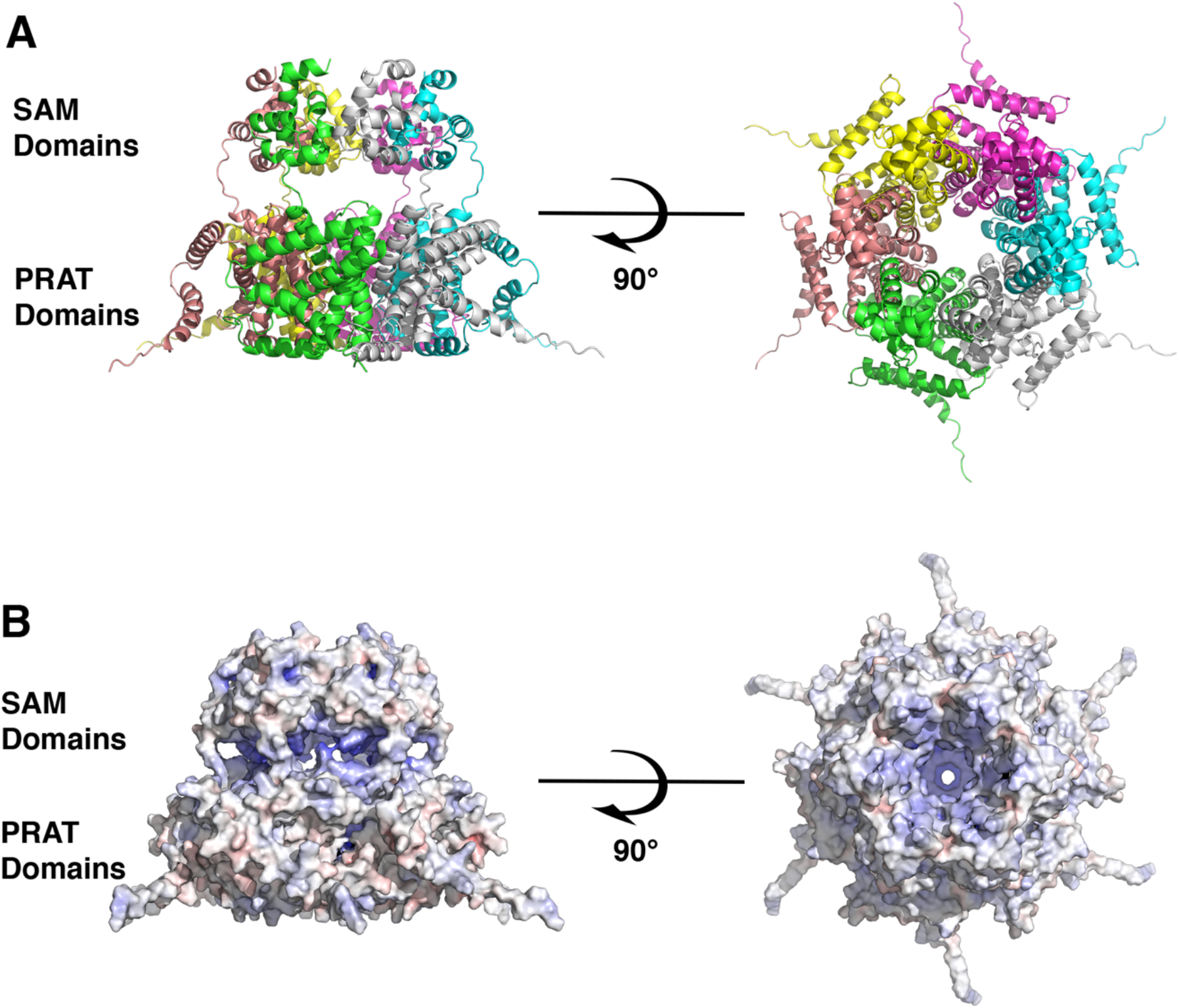
AlphaFold predicted structure of full length Tric1. **A.** A ribbon representation of the predicted structure of the full length Tric1 as a hexamer. Each monomer is shown in a different color. **B.** An electrostatic surface representation of the hexamer. The left panels show the side view of the hexamer and the right panels show the top view from the side containing the SAM domains. Regions of positive potential are colored in blue and regions of negative potential are colored in red.

### Structure predictions of the SAM domain and Tric1 support an oligomeric arrangement for tRNA import

It is unlikely that the SAM domain will adopt a superhelical structure as observed in the crystal structure in the context of the full-length structure which includes the PRAT domain, anchoring the protein in the mitochondrial membrane, however one might expect that a closed cyclic assembly could occur in the presence of the transmembrane domain.

To assess the possible oligomeric arrangement of the protein structure, predictions were performed. The artificial intelligence program, AlphaFold 2, developed by Deep Mind, is a powerful tool for predicting tertiary structures of proteins (32,33). In recent years it has been shown to be remarkably accurate at protein structure prediction, and more recently algorithms have been developed to also predict limited multimeric structures.

Using AlphaFold 2 (version 2.3.1) (34) on the Galaxy Australia web based platform (https://usegalaxy.org.au/), structure predictions of the SAM domain (residues 191 – 261) were carried out as a monomer as well as different multimers (trimer, tetramer, pentamer, hexamer and heptamer). The model confidence scores of the top ranked model for each predicted structure as well as the average pLDDT scores for the top ranked structures are listed in Table S1 and each of the predicted structures is shown in Fig. S5A-E.

The results give high model confidence scores for the trimer, tetramer, pentamer and hexamer as well as high pLDDT scores for each of the top ranked multimeric structures. In contrast the heptamer exhibited a very low model confidence score and a significantly lower pLDDT score. Only in the case of the pentamer and the hexamer was an intact cyclic overall structure able to be predicted (Fig. S5C-D). The trimer and the tetramer formed partially cyclic structures which were superimposable on a portion of the hexamer (Fig. S5C). In all predicted structures, the side chains of Lys234, Asp235 and Arg243 are involved in electrostatic contacts as seen in the crystal structure.

Further AlphaFold 2 predictions were also carried out for the full length Tric1 incorporating both the SAM and the PRAT domains (Fig. S6A). This showed structural conservation for the experimentally determined SAM domain as well as a possible fold for the PRAT domain. The two domains are connected by an extended linker region which exhibits lower pLDDT scores. The pLDDT score for the Tric1 monomer was overall lower than for the SAM domain alone (Table S1) however the scores for the four predicted transmembrane helices of the PRAT domain (residues 55 - 79 and 106 - 127, 137 – 154 and 162 - 186) and the residues of the SAM domain (residues 196 – 261) are considerably higher (>85) than the N terminal helix (1–31), the interdomain linker (185–195) and a predicted helical insertion between the first and second TM helices (residues 80-105) (Fig. S6A). Thus, the prediction gives a good level of confidence for the core regions of the Tric1 structure, with lower predictions, particularly for the N and C termini and the linker regions between the PRAT and SAM domains.

Further predictions were carried out for different multimers of the full length Tric1 with scores for the predictions shown in Table S1. The results of these predictions showed high pLDDT scores (>80) for the SAM domains in all cases. While the scores for the PRAT domains were considerably lower, as the oligomer increased to a pentamer (Fig. S6B) and a hexamer (Fig. S6C) the model confidence scores increased and the pLDDT scores increased, particularly for residues making up the last two transmembrane helices of the PRAT domain (residues 137-154 and 162-186). These predictions showed that a closed oligomer could be formed for the pentamer and hexamer although the exact arrangement of the PRAT domains within the oligomer could not be predicted with high confidence (Fig. S6B, C). This may be due to the ten-residue extended linker sequence separating the PRAT and the SAM domains for each monomer. Interestingly, from the predictions obtained, a cyclic pentameric structure results in a pore through the centre of the oligomer that is approximately 13 Å wide, at the position of the SAM domains (Fig. S7A). In the case of the hexameric structure the pore has a width of approximately 20Å (Fig. S7B). In both cases, the electrostatic nature of the pore shows a positively charged surface throughout, supporting an expected pathway for a negatively charged tRNA molecule to pass through the oligomer, and presumably across the mitochondrial membrane (Fig. S7). Furthermore, the width of an unfolded but still helical tRNA molecule is ∼18 Å supporting the hypothesis that the tRNA is able to thread through the pore of a hexamer.

Based on the structural results observed for the SAM domain as well as the AlphaFold 2 predictions for both the SAM domain and for the full-length protein, we hypothesize that, while a pentameric quaternary structure cannot be ruled out, Tric1 likely forms a hexameric ring structure to facilitate movement of tRNA across the membrane through a pore in the oligomeric assembly.

## Discussion

The mitochondrial outer membrane proteins, Tric1 and Tric2, contain a C-terminal SAM domain that is required for the efficient uptake of tRNA into plant mitochondria (5). Conserved lysine residues within the SAM domain were identified to be essential for tRNA binding ability *in vitro* (5). Inactivation of both Tric1 and Tric2 in Arabidopsis (*tric1:tric2*) resulted in severely chlorotic and developmentally delayed plants, defective in tRNA import ability that could not be restored by complementation using the Tric1 protein lacking the SAM domain, suggesting an essential role.

To further characterise the structural and functional properties of the Tric1/2 SAM domain, we determined its crystal structure to 1.5 Å resolution. The crystal lattice reveals a helical superstructure of 6 monomers per helical turn. Furthermore, solution studies indicate an oligomeric species that is highly thermally stable, suggesting that the oligomerization behaviour of the protein is not a result of crystallization but rather a natural characteristic of the SAM domain. This self-oligomerization is typical of other SAM domain containing proteins and has been implicated in a wide range of protein functions such as mediation of protein-protein interactions, signalling cascades, transcription and DNA repair activities, often associated with the mode of oligomerization (31,35-39).

Mutational and structural studies indicate that Gly241 was required to maintain the hexameric helical structure and not Asp235. EMSA assays, carried out using Asp235Ala and Gly241Glu variants, reveal, in both cases, abolished RNA binding capability. This suggests that the oligomerization ability of the SAM domain is required for tRNA binding, and that removal of the salt-bridge between adjacent SAM domains can also alter RNA binding ability.

To assess the function of both residues in planta, complementation assays were carried out. The double mutant variant (Asp235Ala:Gly241Glu) of the full-length Tric1 was tested and failed to restore the aberrant growth phenotype of the double knock-out line *tric1:tric2*, unlike the full-length WT Tric1. This suggests that these residues are required for functionality of Tric1.

The protein residues previously indicated to be involved in RNA binding (Lys205 and Lys210) (5) reside on the exterior surface of each monomer in the superhelical structure. An inspection of the electrostatic characteristics of the oligomeric structure shows the outer surface as more positively charged. Additionally, each hexameric ring exhibits a charge polarity, with the upper surface of the ring (C terminal end) more negatively charged and the lower surface (N terminal end) more positively charged. It should be noted however that the final three residues at the C terminal end of the domain (Lys259, Arg260, Lys261) are not modelled in the structure as there was insufficient electron density. This may suggest that the C terminal surface is more positively charged than appears in the structure. The N terminus of the SAM domain lies along the outer and bottom (positively charged) face of the hexameric ring. In the full length Tric1/2, the PRAT domain, containing 4 predicted transmembrane alpha helices, would be expected to be positioned near to the N terminal end of the SAM domain.

SAM domains have been shown to form circular oligomeric structures that are required for protein function (40). For example, in SARM1 (sterile alpha and TIR motif containing 1), a human NAD^+^ hydrolase, an octameric assembly is mediated via two tandem SAM domains allowing the formation of an antiparallel double stranded assembly of a C-terminal TIR (Toll/interleukin-1 receptor) domains (41). Guided by these observations, structure predictions were carried out for both the SAM domain as well as the full length Tric1 as different oligomers. It is apparent from these results that the protein can arrange as a cyclic pentamer or hexamer.

Based on the structural properties of the Tric1 SAM domain and the relative position of the predicted transmembrane regions of the full-length protein, we suggest that Tric1 adopts a hexameric ring structure which generates a pore through which tRNA is able to pass to cross the mitochondrial membranes. To validate this hypothesis further, structural and biochemical studies need to be carried out both *in vivo* and *in vitro* with the full-length protein.

## Experimental Procedures

### Clones

Cloning of the full length Tric1 cDNA (At3g49560) and the Tric1 SAM domain (residues 191 – 261) is described previously (5). Mutations corresponding to Asp235Ala, Gly241Glu and Asp235Ala:Gly241Glu were introduced into the previously published vectors (5) using the QuickChange II Site-directed mutagenesis kit (Agilent) using primers listed in Table S2.

### Protein expression and purification

The wildtype (WT) Tric1 SAM domain (residues 191 – 261) and the variants Asp235Ala, Gly241Glu and Asp235Ala:Glu241Gly were recombinantly expressed in *Escherichia coli* Rosetta2 (DE3) using Lysogeny broth (LB) containing 50 μg/ml ampicillin and 34 μg/ml chloramphenicol. Protein expression was induced by the addition of 200 μM Isopropyl β-d-1-thiogalactopyranoside (IPTG) to the growth media and incubated at 20°C for 4 h. Cells were lysed using an EmulsiFlex-C5 high-pressure homogenizer (Avestin) in buffer A (20 mM HEPES, pH 7.5, 50 mM NaCl, 20 mM imidazole). The cell lysate was centrifuged at 24,000 x g for 45 min and the supernatant applied to 2.5 ml of NiNTA beads pre-equilibrated with buffer A. The beads were washed with 20 ml of buffer A, followed by 20 ml of buffer B (20 mM HEPES, pH 7.5, 50 mM NaCl, 60 mM imidazole) and the target protein eluted with 10 ml of buffer C (20 mM HEPES, pH 7.5, 50 mM NaCl, 250 mM imidazole). The fractions containing the protein of interest were applied to a Sephacryl 300 16/60 gel filtration column (GE Healthcare) and eluted with buffer D (10 mM HEPES, pH 7.5, 50 mM NaCl). Protein elution was monitored by absorbance at 280 nm. The purified protein was concentrated to a final concentration of 8-12 mg/ml for crystallization experiments.

### SeMet protein production and purification

A selenomethionine (SeMet) variant of the Tric1 SAM domain was recombinantly expressed using the methionine inhibition pathway approach (42,43) in *E. coli* BL21 (DE3). Seven millilitres of inoculant grown overnight from a single transformed colony was added to M9 minimal media (500 ml) supplemented with 1 mM magnesium sulphate, 0.2% (w/v) glucose 10^-5^% thiamine, 0.25 ml of 0.5 mM calcium chloride, and 50 µg/ml ampicillin. Amino acids were added to the minimal media containing the bacteria to a final concentration of 125 mg/L for lysine, 100 mg/L for phenylalanine and threonine, 50 mg/L for isoleucine, leucine and valine and 60 m/L for L-selenomethionine. Cells were incubated for 1 h at 37°C, protein expression was induced by the addition of 0.4 mM IPTG, for 6 h at 20°C and the cells were then cooled to 4°C for 30 min. The bacteria were harvested and lysed as described above. The SeMet variant protein was purified by NiNTA affinity chromatography using Buffers A and C above supplemented with 10 mM MgCl_2_ and 10 mM βME. The eluted protein was dialysed into 10 mM HEPES pH 7.5, 50 mM NaCl, 1 mM βME and concentrated to 12 mg/ml final concentration for crystallization.

### Analysis of oligomeric formation in solution

To assess formation of oligomers in solution, 1 ml of purified WT SAM domain at 9.5 mg/ml was loaded onto a Superdex 75 16/60 gel filtration column (GE Healthcare) at 1 ml/min pre-equilibrated in buffer D. Samples of the eluted peaks were mixed with 5 X Laemmli sample buffer (250 mM Tris-HCl pH 6.8, 4 %(w/v) sodium dodecyl sulfate (SDS), 30 %(v/v) glycerol, 0.6 g/L bromophenol blue). 3 μl of each sample was heated at 95 °C for 10 minutes and loaded onto a 15 %(w/v) SDS-PAGE gel alongside 3 µl of the same sample that had not been heated, and 5 µl of prestained protein ladder (New England Biolabs). The gel was electrophoresed at 150 V, 35 mA until the dye front was 0.5 cm from the base of the gel. The gel was then blotted onto a nitrocellulose membrane (GE Healthcare) using the XCell II blot module (Thermo Fisher) at a constant current of 170 mA for 1 h at 4 °C. The membrane was incubated in 20 ml blocking buffer (50 mM Tris-HCl pH 7.5, 150 mM NaCl, 0.05 %(v/v) TWEEN 20, 5 %(w/v) skim milk powder) for 1 h at RT with gentle shaking. A monoclonal anti-polyhistidine-peroxidase antibody was added at a 1/2000 dilution (Sigma-Aldrich). The membrane was incubated with the antibody for 10 min at RT with gentle shaking. The antibody solution was removed and the membrane washed 5 x with tris buffered saline with tween (TBST) (50 mM Tris-HCl pH 7.5, 150 mM NaCl, 0.05 %(v/v) TWEEN 20) for 10 minutes each with gentle shaking at RT. The membrane was removed from the TBST and 1 ml of WesternSure chemiluminescent substrate (LI-COR) was pipetted over the surface. The blot was imaged using a chemidoc imaging system (BioRad).

### Crystallization

Crystallization of the Tric1 SAM domain (WT) and variants were carried out by vapor diffusion using the hanging-drop method at 20°C. Rod shaped crystals of the WT and Asp235Ala were obtained using a protein concentration of 8 - 10 mg/ml and a precipitant solution of 1.5 - 2.0 M ammonium sulfate, 0.1 M Tris, pH 8.0 - 8.7. Microcrystals of the Gly241Glu mutant were obtained using protein at 16 mg/ml and a precipitant solution of 1.8 - 2.3 M ammonium sulfate, 0.1M Tris, pH 8.0 - 8.7. Crystals of suitable size for diffraction experiments were obtained through a microseeding procedure and also by gradually increasing the precipitant concentration in the reservoir solution to a final ammonium sulfate concentration of 2.0 M.

Crystals of the SeMet mutant protein were obtained by the sitting-drop vapor diffusion method at 20 °C using protein at 6 - 12 mg/ml and a precipitant solution of 0.8 - 1.0 M ammonium sulfate, 0.1 M Tris, pH 8.0 - 8.5.

### X-ray data collection and data processing

X-ray diffraction data was collected at the Australian Synchrotron beamlines MX1 and MX2 equipped with an EIGER 9M detector and an EIGER 16M pixel detector respectively with continuous readout (‘shutterless’ data collection). All data was collected from 360° of crystal rotation using the Blu-Ice data acquisition software (44), integrated using the XDS processing package (45) and data reduction carried out using the CCP4 software package (46). Phasing of the SeMet mutant was carried out using the multi anomalous diffraction method from selenium atoms in the mutant protein. The structures of the Asp235Ala and Gly241Glu variant proteins were solved by molecular replacement with the program PHASER (46) using the WT Tric1 SAM domain as the search model. Iterative cycles of model building using the graphic software COOT (47) and crystallographic refinement using the PHEINX software suite (48) was carried out. The data reduction and refinement statistics are given in Table 1.

### Protein structure prediction

Artificial intelligence guided 3D protein structure predictions were carried out using AlphaFold version 2.3.1 (34) on the Galaxy Australia web based platform (https://usegalaxy.org.au/). Predictions were carried out on the Tric1 SAM domain alone (residues 191 - 261) and the full length Tric1 as monomers. Predictions of multimers were carried out using AlphaFold in ‘Multimer” mode where multiple sequences (depending on the oligomeric state being predicted) were input in FASTA format. All predictions were carried out using the full database for enhanced accuracy.

### Electrophoretic mobility shift assay

RNA electromobility shift assays were performed using 5’-(6-FAM)-labelled RNA oligonucleotides corresponding to the T-arm of tRNA^ala^ (Table S1). Fluorescein-labelled tRNA^ala^ oligonucleotides (120 nM) were incubated with 160 µM purified protein in a binding buffer (15 mM Tris-HCl pH 7.0, 150 mM NaCl, 10% (w/v) glycerol) and 2 units/μL RNase inhibitor RiboShield (PCR Biosystems) for 15 minutes at 22°C. The reaction was resolved on a nondenaturing 6% Tris/borate/EDTA polyacrylamide gel, in 0.5X Tris/borate/EDTA buffer gel and imaged using an Amersham Typhoon scanner (GE Healthcare).

### Complementation of *tric1:tric2*

Complementation of the *tric1:tric2* double mutant (5) was carried out using *Agrobacterium tumefaciens* mediated transformation.. Two independent T_3_ complementation lines were used for all analysis. Chlorophyll content was determined using the SPAD-502 chlorophyll meter (Minolta) and averaged from 10 measurements taken from distinct rosette leaf areas with five individual plants.

## Data Availability

The X-ray crystal structure data presented in this study have been deposited to the Worldwide Protein Data Bank. The PDB accession codes for the WT, D235A and G241E Tric1 SAM domains are 8UCY, 8UCZ and 8UD0 respectively.

## Supporting Information

This article contains supporting information.

## CRediT Author Statement

**Bence Olasz:** Investigation, Validation, Formal analysis, Writing – Original Draft. **Luke Smithers:** Investigation, Validation, Writing – Review and Editing. **Genevieve Evans:** Conceptualization, Methodology, Investigation. **Anandhi Anandan:** Investigation. **Monika Murcha:** Conceptualization, Methodology, Validation, Formal analysis, Investigation, Writing – Review and Editing, Supervision. **Alice Vrielink:** Conceptualization, Project administration, Supervision, Methodology, Investigation, Writing-Reviewing and Editing.

## Abbreviations

SAM: (sterile alpha motif)
Tric: (tRNA import component)
PRAT: (preprotein and amino acid transporters)

## Acknowledgments

B.O acknowledges the Commonwealth Governments support through the University Postgraduate Award and Australian Government Research Training Program Scholarships at The University of Western Australia. This research was undertaken on the MX1 and MX2 beamlines at the Australian Synchrotron, part of the Australian Nuclear Science and Technology Organisation (ANSTO) and made use of the Australian Cancer Research Foundation (ACRF) detector. The authors thank Dr. Daniel Eriksson at the Australian Synchrotron for assistance with X-ray data collection and Maja Tjørnelund Jensen for assistance with protein production.

