## Supplementary Figures and Tables for "Structural analysis of the Sterile alpha motif (SAM) domain of the Arabidopsis mitochondrial tRNA import receptor"

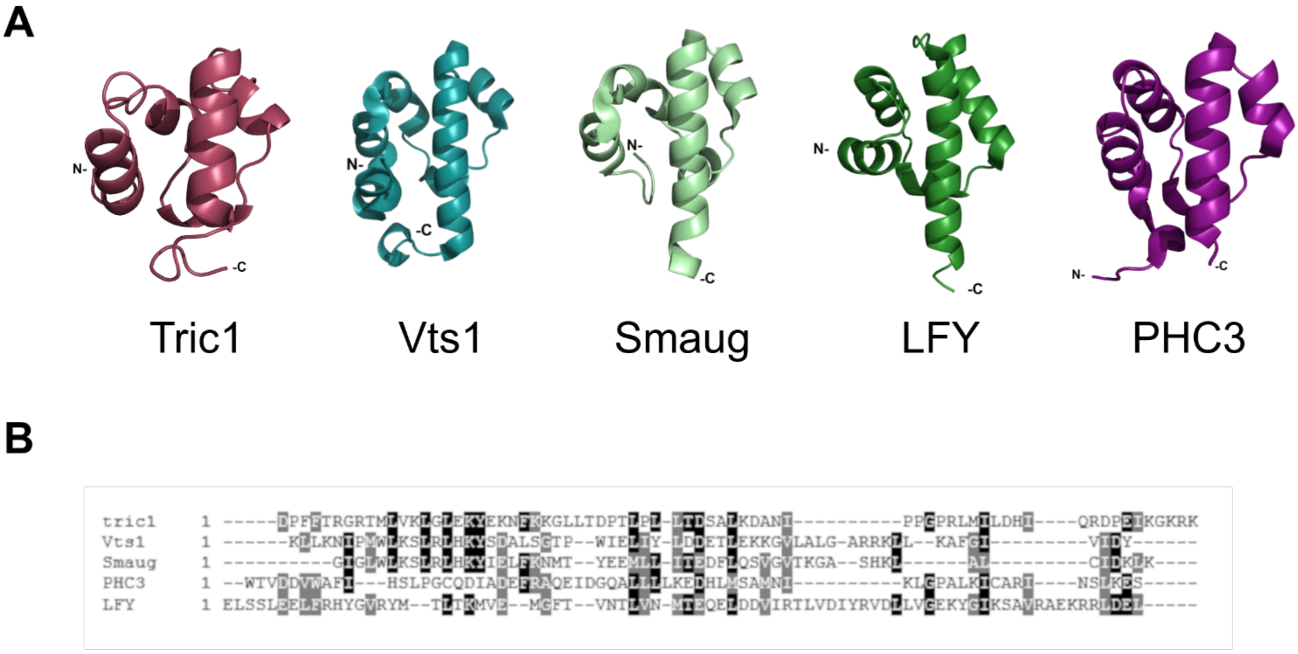


**Supplementary Figure 1: Comparisons of SAM domain structure from different proteins. A. Tric1:** plant RNA binding protein involved in the transport of tRNA into the mitochondria. Vts1: yeast RNA binding protein involved in post transcriptional regulation (PDB code 2FE9). Smaug: *Drosophila melanogaster* RNA binding protein required for abdominal segmentation in early embryos (PDB code 1OXJ). LFY: plant SAM domain containing protein involved in DNA binding regulating floral transition (PDB code: 4UDE). PHC3: human protein required to maintain the transcriptionally repressive state of many genes throughout development (PDB code 4PZO). **B.** Sequence alignment of the SAM domains from each of the protein displayed in panel A).

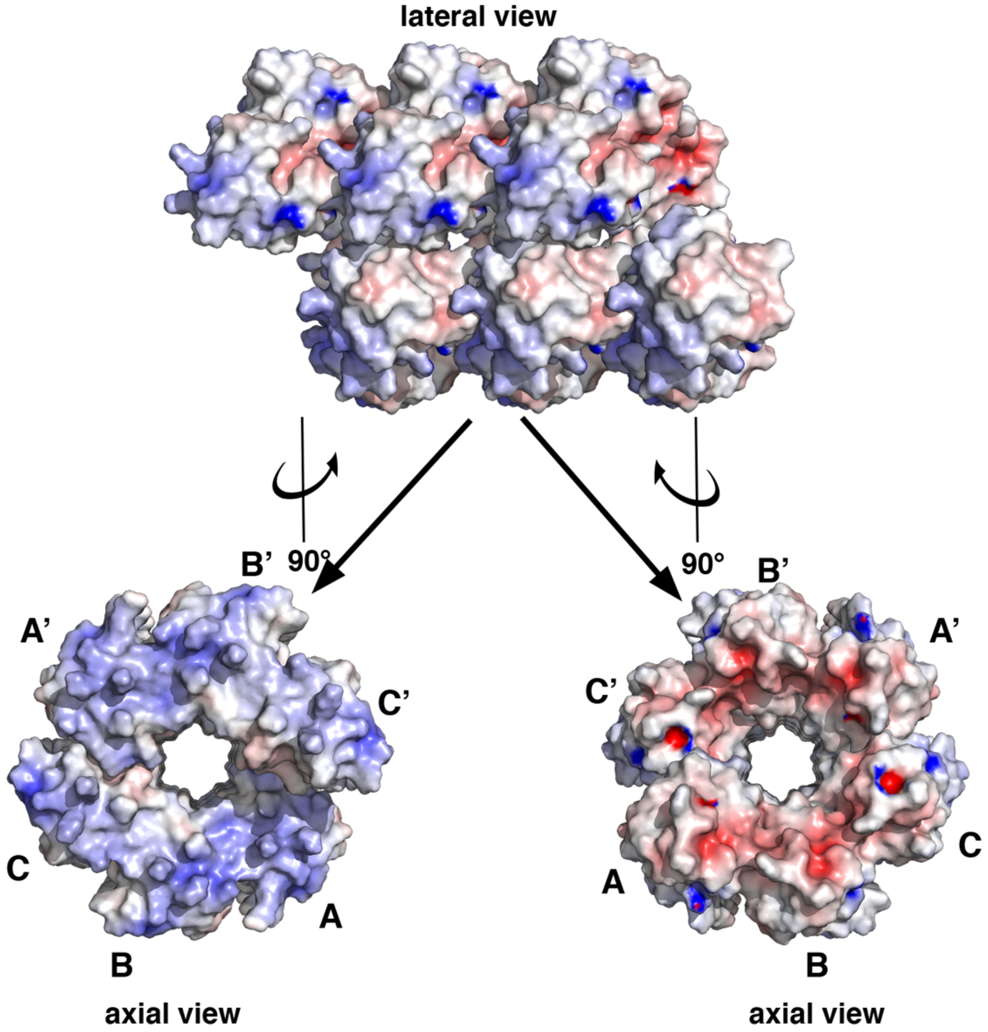


**Supplementary Figure 2: Electrostatic potential surface representation of the SAM domain**. The top panel shows the lateral view consisting of 18 monomers of the SAM domain. The bottom two panels shows the two axial views corresponding to 90 ° rotations of the helical superstructure. The monomers in a single asymmetric unit of the crystal structure are indicated by A, B and C and those in a second asymmetric unit are labelled as A’, B’ and C’. Regions of positive potential are colored in blue and regions of negative potential are colored in red.

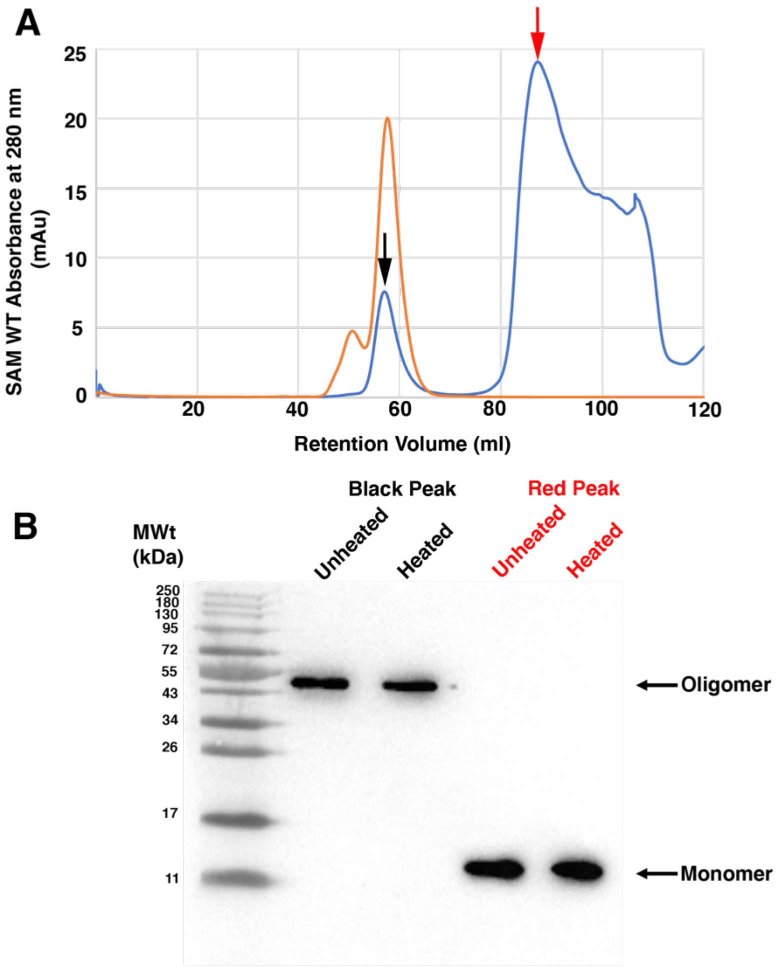


**Supplementary Figure 3: Analysis of SAM domain oligomerization in solution. A).** Chromatogram obtained by size exclusion chromatography of SAM domain from a Superdex S75 16/60 gel filtration column. The blue trace represents the chromatogram of the SAM domain and the orange trace represents the chromatogram of a BSA standard. **B).** Western blot analysis of the indicated fractions obtained from the size exclusion chromatogram shown in A. The samples were electrophoresed on a 15% SDS-PAGE gel and blotted onto a nitrocellulose membrane which was then treated with a monoclonal anti-polyhistidine-peroxidase antibody. Samples for each peak were applied to gel unheated as well as heat treated for 10 minutes at 95°C. The bands corresponding to the WT SAM domain monomer and oligomer are labelled with red and black arrows respectively.

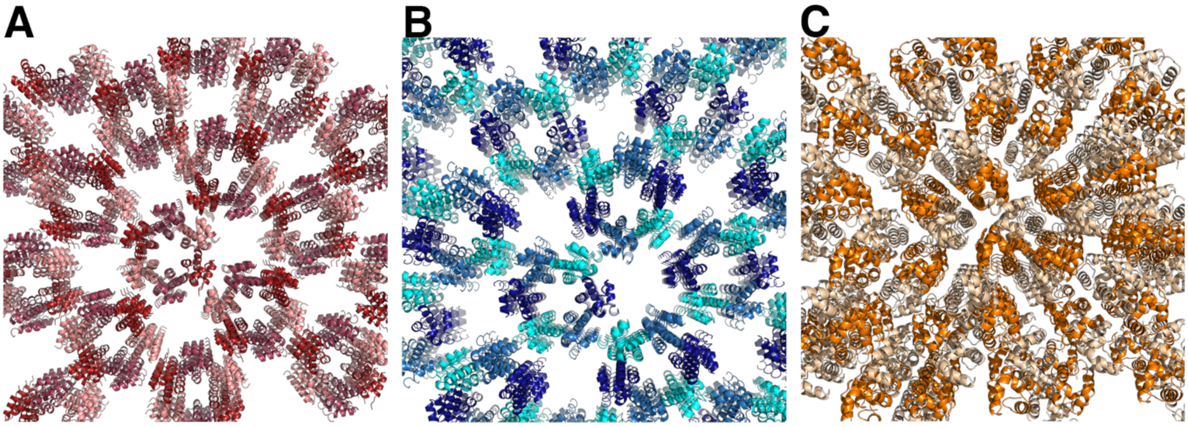


**Supplementary Figure 4: The crystallographic lattice symmetry of WT Tric1 SAM domain and mutant structures. A.** The WT protein lattice **B.** The lattice of the Asp235Ala mutant **C.** the lattice of the Gly241Glu mutant. The monomers in the asymmetric units are colored as shown in Figures 1 and 2.

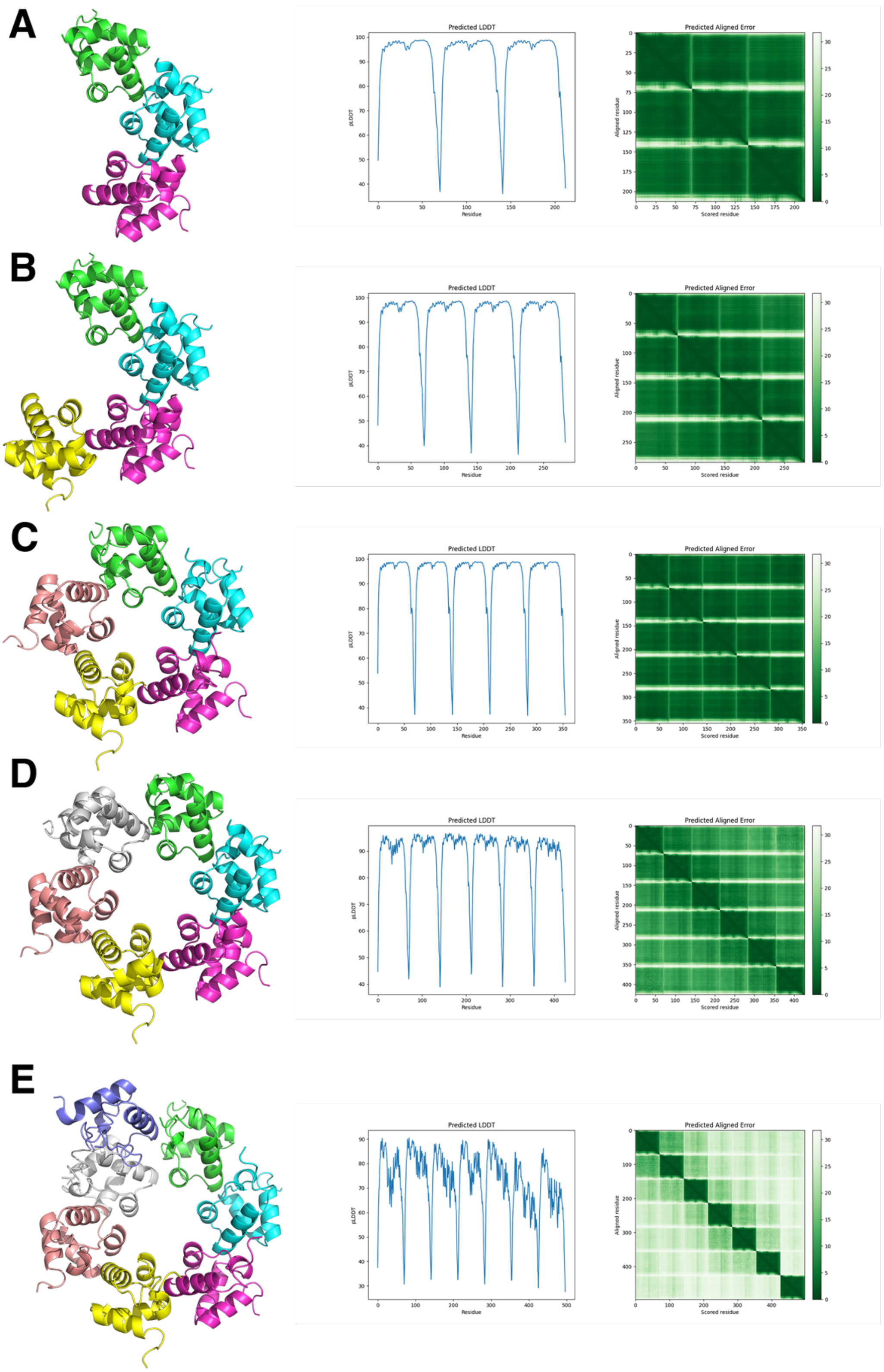


**Supplementary Figure 5: AlphaFold predicted structures of the Tric1 SAM domain (residues 191-261).** A ribbon representation of the predicted structures as **A.** a trimer, **B.** a tetramer, **C.** a pentamer, **D.** a hexamer and **E.** a heptamer. The pLDDT scores and the predicted alignment error analyses as a function of residues as output from AlphaFold are shown to the right of each predicted structure. Each monomer in the multimeric structures are shown in a different color.

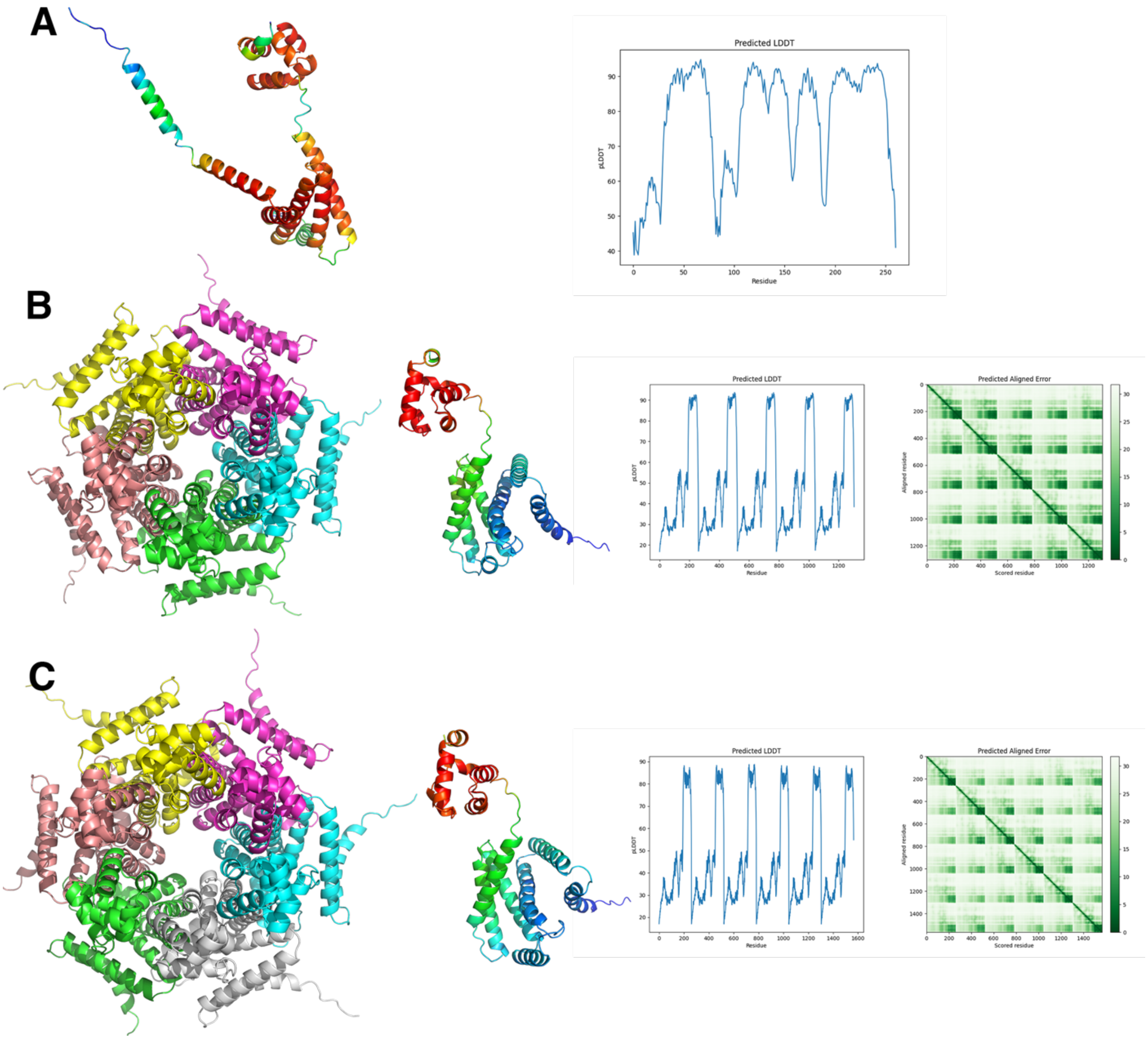


**Supplementary Figure 6: AlphaFold predicted structures of full length Tric1.** A ribbon representation of the predicted structures as **A.** a monomer, **B.** a pentamer and **C.** a hexamer. For panel A the ribbon representations is colored by pLDDT score (with increasing score from blue to red). For panels B and C the left ribbon diagram shows the multimer colored by chain and the middle panel shows the monomer, colored by pLDDT score (with increasing score from blue to red). The right panels show the pLDDT scores and the predicted alignment error analyses as a function of residues as output from AlphaFold.

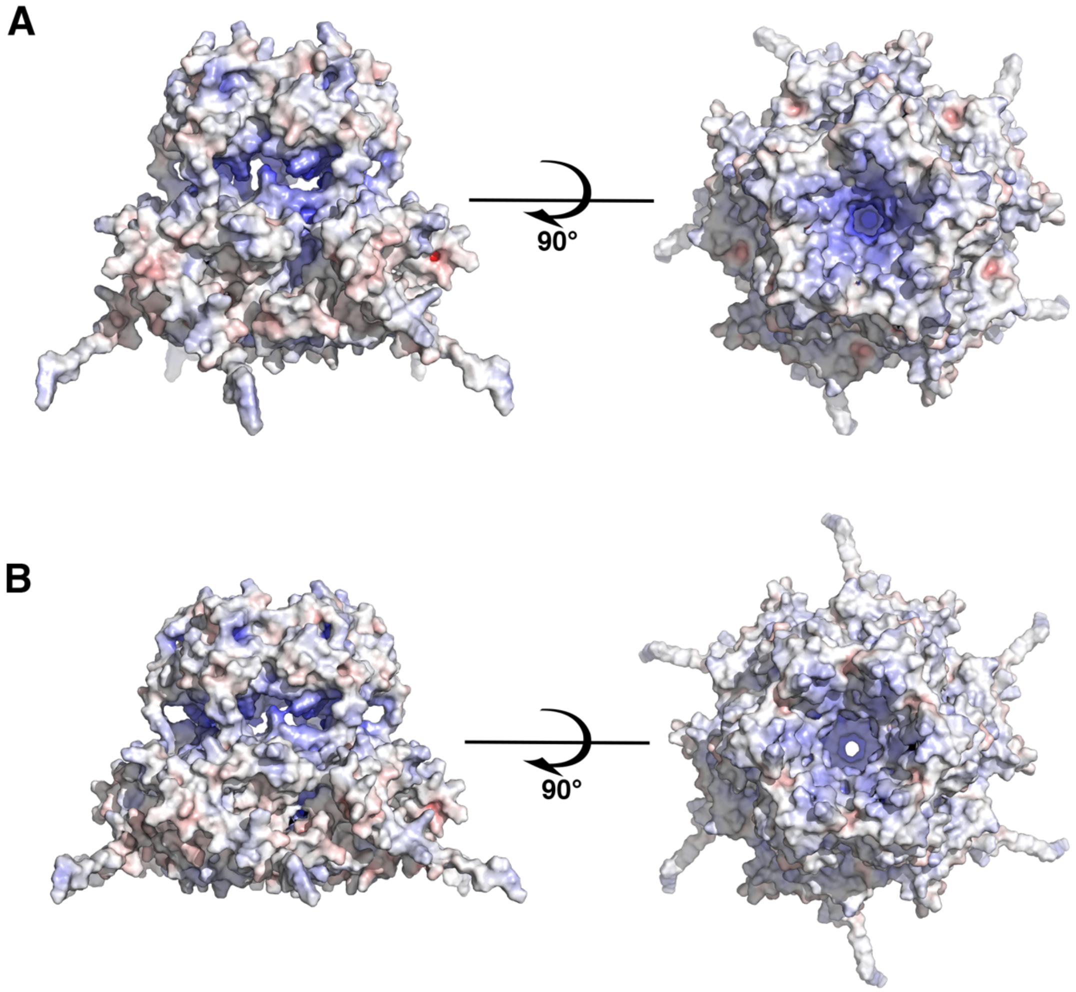


**Supplementary Figure 7: Electrostatic surfaces of full length Tric1 pentamer and hexamer.** A surface representation of the AlphaFold predicted oligomers of **A.** a pentamer and **B.** a hexamer. Each panel is shown in 2 different views. Regions of positive potential are colored in blue and regions of negative potential are colored in red.

**Supplementary Table 1: Structure prediction scores obtained from AlphaFold calculations on SAM domain and on full length Tric1**

| Multimer | Tric1 SAM Domain (Model Confidence Score/Average residue pLDDT score for top ranked structure) | Full length Tric1 (Model Confidence Score/Average residue pLDDT score for top ranked structure) |
|---|---|---|
| Monomer | 91.87/91.4 | 78.29/78.7 |
| Trimer | 0.86/91.0 | 0.33/45.8 |
| Tetramer | 0.84/90.6 | 0.27/38.7 |
| Pentamer | 0.90/91.8 | 0.52/50.5 |
| Hexamer | 0.71/87.5 | 0.43/48.4 |
| Heptamer | 0.29/72.4 | Not determined |

**Supplementary Table 2 – Primers and RNA sequences**

| | | |
|---|---|---|
| **Tric1** | SDM primer for Tric1 Asp234Ala | F: TAGCGCGCTGAAAGCTGCGAACATCCC |
| | | R: GGGATGTTCGCAGCTTTCAGCGCGCTA |
| **Tric1** | SDM primer for Tric1 Gly241Glu | F: GAACATCCCACCAGAGCCAAGACTTATGATAC |
| | | R: GTATCATAAGTCTTGGCTCTGGTGGGATGTTC |
| **tRNA$^{ala}$-T arm** | RNA binding | 5’-(6FAM)-UCGCUUUGCAUGCGA |
